# Unraveling the implications of environmental, host and pathogen-specific determinants on clinical presentations and disease progression among Indian melioidosis patients

**DOI:** 10.1101/463687

**Authors:** Tushar Shaw, Chaitanya Tellapragada, Asha Kamath, Vandana Kalwaje Eshwara, Chiranjay Mukhopadhyay

## Abstract

**Background:** Melioidosis is gaining recognition as an emerging infectious disease with diverse clinical manifestations and high-case fatality rates,worldwide. However, the molecular epidemiology of the disease outside the endemic regions such as,Thailand and Northern Australia remains unclear.

**Methods:** Clinical data and *B. pseudomallei* (Bps) isolates obtained from 199 culture-confirmed cases of melioidosis, diagnosed during 2006-2016 inSouth India were used to elucidate the host and pathogen-specific variable virulence determinants associated with clinical presentations and disease progression. Further, we determined the temporal variations and the influence of ecological factors on Bps Lipopolysaccharide (LPS) genotypes causing infections.

**Result:** Severe forms of the disease were observed amongst 169 (85%) patients. Renal dysfunction and infection due to Bps harboring Bim-ABm variant had significant associations with severe forms of the disease. Diabetes mellitus, septicemic melioidosis and infection due to LPS-B genotype were independent risk factors for mortality. LPS-B (74%) and LPS-A (20.6%) were the prevalent genotypes causing infections. Both genotypes demonstrated temporal variations and had significant correlations with rainfall and humidity.

**Conclusion:** Our study findings suggest that the pathogen-specific virulence traits, under the influence of ecological factors are the key drivers for geographical variations in the molecular epidemiology of melioidosis.

## Introduction

Melioidosis is a fatal infectious disease caused by soil saprophytic bacteria, *Burkholderia pseudomallei.* Infection occurs mostly through the inhalation or percutaneous inoculation of the bacteria from contaminated soil or surface water. The disease manifests with diverse clinical presentations ranging from mild localized infection to fulminant sepsis. *B. pseudomallei*, being a soil saprophyte, uses horizontal gene transfers as a mechanism for its persistence both in the environment and the host. It is possible that the virulence attributes of *B. pseudomallei* can significantly vary under the influence of regional environmental/ecological conditions, in turn, leading to the occurrence of distinct clinical manifestations. Lipopolysaccharide (LPS) of *B. pseudomallei*, is a well-known virulence factor that confers serum resistance and helps in evading host immune defenses during the early stages of infection. In this context, LPS is gaining recognition as a potential candidate for vaccine and diagnostic assay development [1,2]. Three genotypes of LPS namely A, B and B2 were reported previously with variations in their geographic distribution and ability to induce immune responses in animal models [3]. Burkholderia intracellular motility (BimA) and filamentous hemagglutinin gene (fha B3) were reported previously as the significant variable virulence factors based on their geographic distribution and associations with clinical presentations [4].

Amidst the ambiguity of its true incidence, melioidosis is gaining importance as an emerging infection in the Indian subcontinent [5-7]. Sporadic cases reported from different parts of the country have shown assorted clinical presentations [7-10]. Distinct/novel sequence types of *B. pseudomallei* using multi-locus sequence typing(MLST) were previously reported from the south-western coastal part of India [11].This geographic region might presumably be one of the endemic hotspots of melioidosis in India reporting several cases [6,7,10]. Clinical isolates of *B. pseudomallei* from this region were genetically diverse from the Australian and Southeast Asian isolates and the prevalent sequence type (ST 1368) lacked significant association with any particular clinical presentation of the disease [11]. With this background, the present study was undertaken: 1) To study the frequencies of variable virulence factors of *B. pseudomallei* and their association with clinical presentations among Indian melioidosis patients. 2) To study the temporal variations and influence of ecological factors on commonly occurring LPS genotypes of *B. pseudomallei* in the south western coastal part of India. As an additional outcome of the study, we elucidated the host and pathogen-specific determinants for mortality in our settings.

## Material and Methods

### 2a. Study site and population

The present study was carried out at a university and referral teaching hospital (2030 bedded) catering to residents residing in Udupi and the surrounding regions of south-western coastal parts of Karnataka, India covering nearly 150-200 km radius of geographical area. The study region experiences tropical climatic conditions with an annual rainfall of >4000 mm during the months of June-October. Microbiological culture confirmed cases of melioidosis, reported during a period of ten years (2006-2016), at the study hospital were included in the study. The study was approved by the Institutional Ethical Committee Kasturba Hospital, Manipal.

### 2b. Study Isolates

A total of 199 non-repetitive, *B. pseudomallei* isolates obtained from similar number of patients diagnosed with melioidosis were included. Isolates from non-blood clinical specimens (respiratory, abscess aspirates, wound swabs and pus) were included from only those patients that had localized form of the disease. When bacteremic melioidosis was encountered, only isolates from blood were studied. For the extraction of bacterial DNA, QIAamp DNA mini kit (Qiagen, Hilden, Germany) was used as per the manufacturer’s instructions. Before inclusion, all the study isolates were confirmed as *B. pseudomallei* using a species-specific PCR targeting the TTS1 gene cluster as described previously [12].

### 2c. Detection of LPS genotypes and variable virulence genes

Collectively, we aimed at detection of LPS genotypes (A, B and B2), BimA gene variants (BimA_Bp_and BimA_Bm_) and fhaB3.

#### Detection of LPS genotypes

Multiplex PCR assay designed to simultaneously detect three different LPS genotypes LPS A, B, and B2 was used in the present study as reported previously [3]. The PCR reaction was set at a final volume of 25 μl using JumStart^TM^Taq Ready Mix (Sigma-Aldrich).Amplification was carried out in a Master cycler gradient (Eppendorf, Hamburg, Germany) with an initial denaturation at 95°C for 10 min, followed by 35 cycles of 95°C for 30s, 59°C for 30 sec, 72°C for 30s and a final extension step of 72°C for 7 min. Details regarding the oligonucleotide primers used in the present study and the expected amplicon sizes are tabulated below (**Table 1**).

**Table 1:**
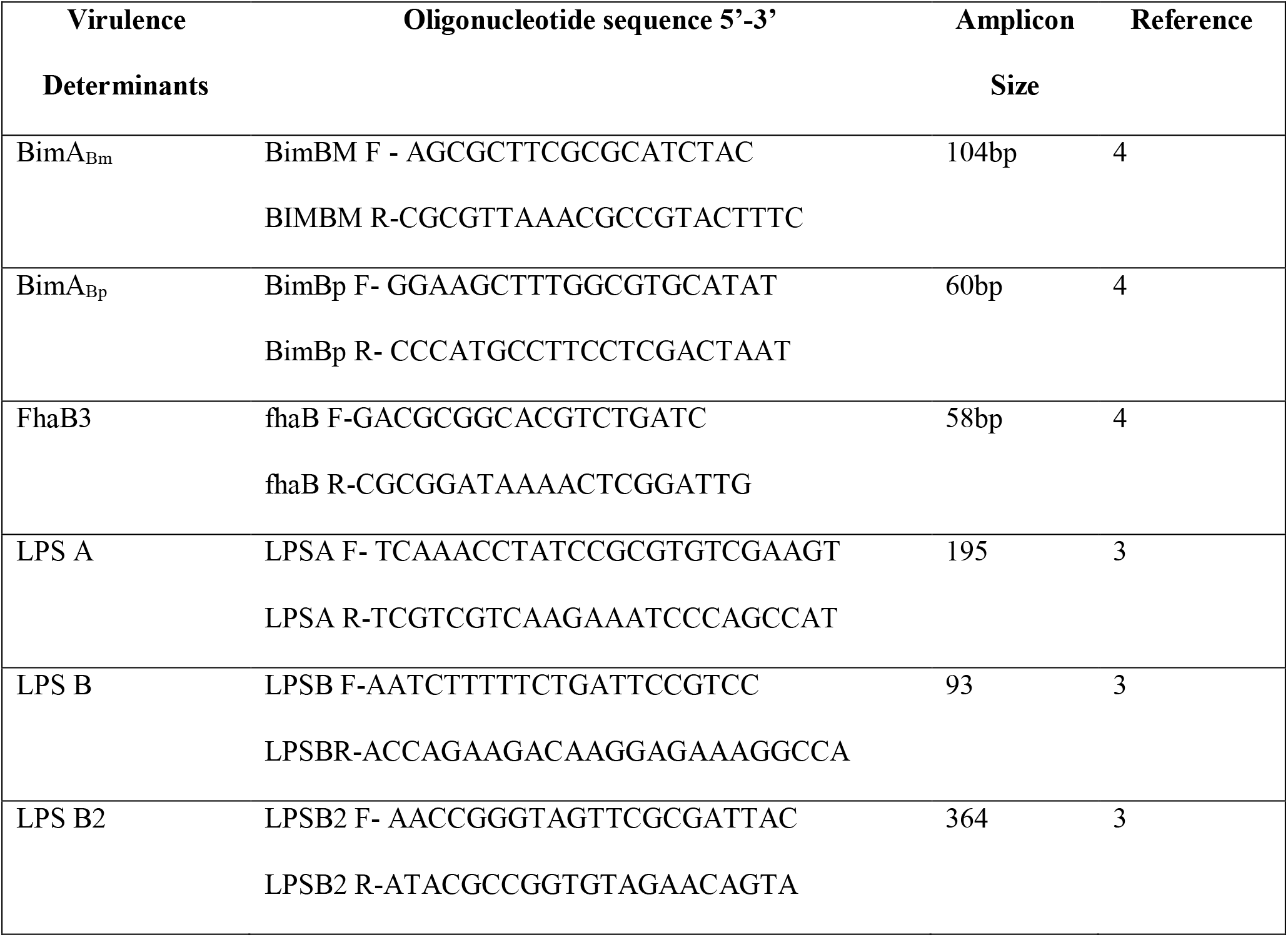
List of oligonucleotide primers used to detect virulence genes of *B. pseudomallei* in the present study.

#### Bim A detection

Both variants of Bim A, namely BimA_Bm_ and BimA_Bp_ were detected using previously reported PCR primers [4]. PCR reaction volumes and cycling conditions for the Bim A genes detection were similar to that of the LPS genotypes detection, except for a change in the annealing temperature to 56^0^C for 1 min. Presence of BimA_Bm_ and BimA_Bp_ were considered when amplicons sized 104 bp and 60 bp respectively were positive on 2.5% Agarose gel stained with 0.5% ethidium bromide.

#### Filamentous hemagglutinin (fha) B3

fha B3 gene was detected using 0.3 uM of each forward and reverse primers to generate a 58 bp product [4]. Amplification was carried using similar cycling conditions as mentioned above for LPS genotypes detection. However, PCR for the detection of fhaB3 was performed separately considering the similar size of the amplicons for both BimA_Bp_ and fha B3.

### 2d. Clinical, epidemiological and meteorological data

Clinical and epidemiological data were extracted in structured study forms. Monthly rainfall and relative humidity data for a period of six years (2010-2016) were obtained from the Indian Meteorological Department, Pune, India.

### 2e. Case definitions

Microbiological culture of blood and/ or other clinically relevant specimens was the mainstay for laboratory diagnosis of melioidosis. For analysis and reporting purposes, the following case definitions were used in the present study:

a) Bacteremic melioidosis: Patients with positive blood cultures for *B. pseudomallei*, with or without culture positivity of other clinically relevant specimens.
b) Pulmonary melioidosis: Patients with clinical and radiological evidence of pneumonia / lower respiratory infection and isolation of *B. pseudomallei* from respiratory specimens and/or blood.
c) Neurological melioidosis: Patients with clinical and radiological evidence of infection affecting brain tissue, meninges or spinal cord and isolation of *B. pseudomallei*from blood, cerebro-spinal fluid or exudates.
d) Septicemic melioidosis: Patients with features of sepsis (fever/hypothermia, leukocytosis, hypotension, pulse rate >90 beats/min, respiratory rate >18/min) and isolation of *B. pseudomallei* from any clinical specimen.
e) Osteo-articular melioidosis: Patients with bone/joint infections such as osteomyelitis and septic arthritis, and isolation of *B. pseudomallei* from any clinical specimen.
f) Localized melioidosis: Patients with skin and soft tissue (or any other single-site) infections that had no bacteremia and radiological or clinical evidence, suggestive of other organ involvement and isolation of *B. pseudomallei* from relevant clinical specimen.

### 2e. Statistical Analysis

Descriptive statistical tools were used to determine the frequencies of categorical study variables. Pearson’s Chi-square test was used to check for the presence of any significant association of host and pathogen characteristics with individual clinical presentations. Risk factors for various clinical presentations and outcomes among the study population were determined using univariate analysis and multivariate logistic regression models (SPSS, version 16). Time series analysis was used to decompose the trend, seasonal and residual components for the climate data and the LPS genotypes. Correlations of the climate data with the LPS genotypes was obtained using Pearson’s correlation coefficient. Generalized additive model was used to predict the LPS genotypes using the climate data. Analysis was carried out using R version 3.3.3. All values were considered significant with p≤0.05.

## Results

### Baseline demography and clinical data

Mean age of our cases was 47.70±15.4 years, with an age range of the study population between 7-86 years. Of the 199 patients, 187 (94%) were from Karnataka and 12 patients (8 from Kerala and 4 from Goa) were from other states on the south-western coastal part of India. Geographical distribution of our study cases is depicted below (Figure 1). Majority of the patients were males (153, 76.9%). 148 (74.3%) and 51 (25.7%) cases were observed during monsoon and dry seasons respectively.

**Figure 1:**
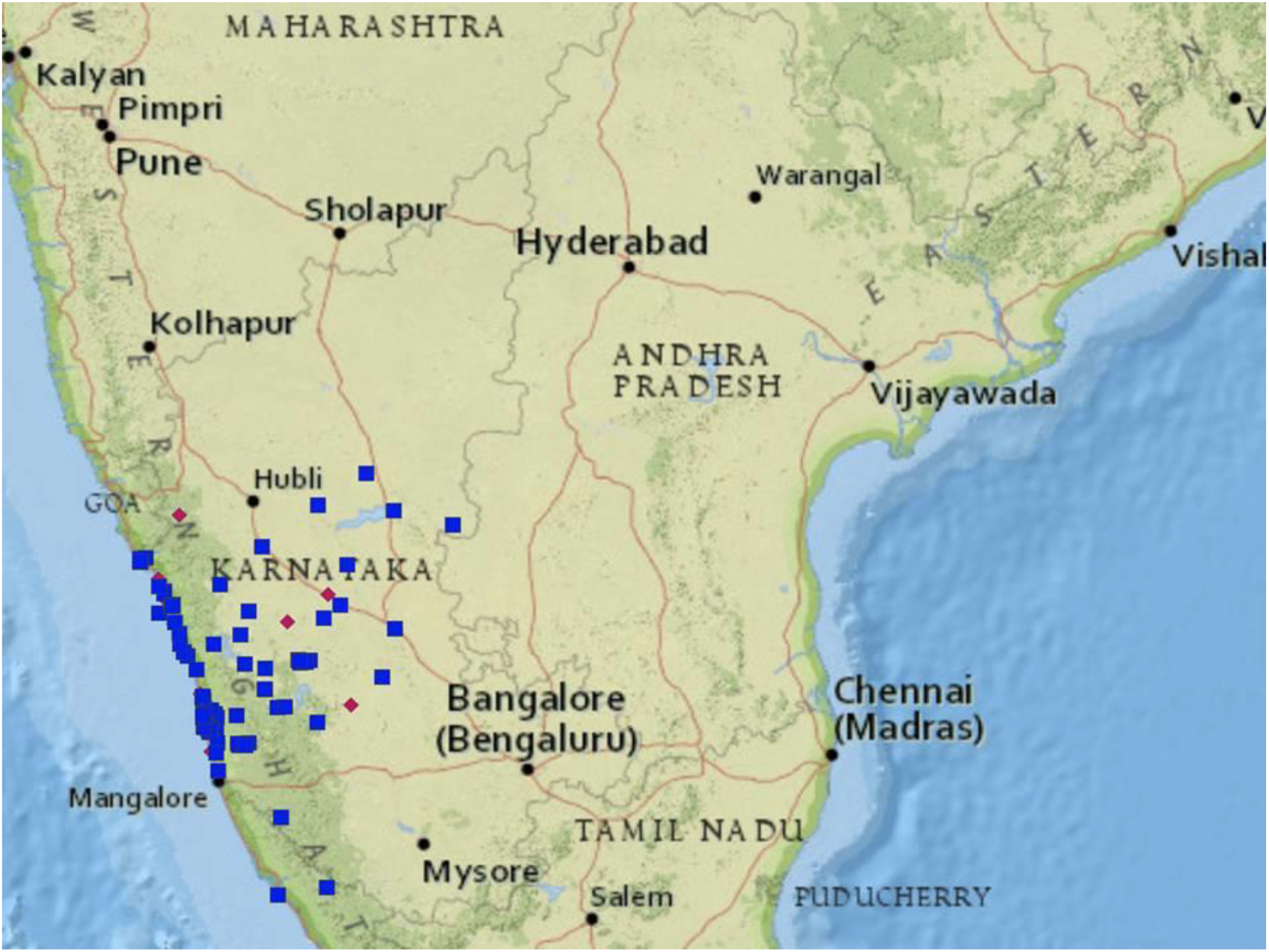
Geographical distribution of study patients and infecting LPS genotypes (Map generated using http://mapmaker.ecdc.europa.eu.). LPS-B genotypes: Blue squares, LPS-A genotypes: Red diamonds.

Of the total 199 cases included in the study, 169 (85%) patients had either one or more of the severe forms of the disease. Severe cases of melioidosis included patients with bacteremic (97, 48.7%), pulmonary (71, 35.6%), septicemic (50, 25.1%) osteo-articualar and/or deep organ abscess (36, 18%) and neurological (22, 11%) forms of the disease. Amongst the 97 cases with bacteremic melioidosis, 47 (48.4%) patients had bacteremia with no other focal infections diagnosed clinically or by radiological examination. Amongst the 50 cases of septicemic melioidosis, 38 (76%) patients had bacteremia and the rest 12 patients had osteo-articular and/or pulmonary forms of the disease without bacteremia. Non-severe or localized form of the disease was observed amongst 30 (15%) patients.

### Association of host factors with clinical presentations

Diabetes mellitus (124, 62.3%) and renal dysfunction (27, 13.5%) were the common co-morbid illnesses observed among the cases. Three patients had malignancy, one patient was a known case of thalassemia and none of the patients were known cases of HIV infection. Associations of individual baseline demographic and clinical characteristics with clinical presentation of the disease among the study population are reported below Table 2. Using univariate analysis, we observed that patients with renal dysfunction had 5.9 (Crude OR: 5.9; 95% CI: 2.86-12.40) and 3.4 (Crude OR: 3.4; 95% CI: 1.50-7.13) times more odds for developing septicemic and bactermic forms of the disease respectively.

**Table 2:** Association of demographical, seasonal and premorbid illnesses with clinical forms of the disease in our study population.

| Variables (N=199) | Bacteremia<br>(n=97) | Pulmonary<br>(n=71) | Neurologic<br>al (n=22) | Sepsis<br>(n=50) | Localized<br>(n=30) | OA & DOA<sup>#</sup><br>(n=36) |
| --- | --- | --- | --- | --- | --- | --- |
| **<u>Age groups in yrs</u>** |  |  |  |  |  |  |
| <18 (14) | 5 (35.7) | 5 (35.7) | 0 | 2 (14.3) | 4 (28.6) | 4 (28.6) |
| 19-50 (99) | 43 (43.4) | 35 (35.4) | 14 (14.1) | 24 (24.2) | 17 (17.2) | 18 (18.2) |
| >51 (85) | 49 (57) | 31 (36) | 8 (9.3) | 24 (27.2) | 9 (10.5) | 14 (16.3) |
| p-value | 0.111 | 0.995 | 0.227 | 0.530 | 0.153 | 0.541 |
| **<u>Gender</u>** |  |  |  |  |  |  |
| Male (153) | 73 (47.7) | 53 (34.6) | 17 (11.1) | 34 (22.2) | 22 (14.4) | 32 (20.9) |
| Female (46) | 24 (52.2) | 18 (39.1) | 5 (10.9) | 16 (34.8) | 8 (17.4) | 4 (8.7) |
| p-value | 0.596 | 0.577 | 0.963 | 0.085 | 0.617 | **0.059** |
| **<u>Season</u>** |  |  |  |  |  |  |
| Monsoon (148) | 78 (52.7) | 56 (37.8) | 17 (11.5) | 41 (27.7) | 22 (14.9) | 23 (15.5) |
| Dry (51) | 19 (37.3) | 15 (29.4) | 5 (9.8) | 9 (17.6) | 8 (15.7) | 13 (25.5) |
| p-value | 0.057 | 0.279 | 0.741 | 0.153 | 0.888 | 0.111 |
| **<u>Diabetes mellitus</u>** |  |  |  |  |  |  |
| Yes (124) | 61 (49.2) | 49 (39.5) | 14 (11.3) | 36 (29) | 18 (14.5) | 26 (21) |
| No (75) | 36 (48) | 22 (29.3) | 8 (10.7) | 14 (18.7) | 12 (16) | 10 (13.3) |
| p-value | 0.870 | 0.146 | 0.892 | 0.102 | 0.777 | 0.175 |

| **Renal dysfunction** |  |  |  |  |  |  |
| --- | --- | --- | --- | --- | --- | --- |
| Yes (27) | 20 (74.1) | 15 (55.6) | 0 | 18 (66.7) | 3 (11.1) | 5 (18.5) |
| No (172) | 77 (44.8) | 56 (32.6) | 22 (100) | 32 (18.6) | 27 (15.7) | 31 (18) |
| p-value | **0.005** | 0.20 | **0.049** | >0.001 | 0.536 | 0.950 |

#Osteo-articular and Deep Organ Abscess.

### Frequencies of individual virulence factors and their association with clinical presentations

Majority of our study isolates belonged to LPS B (n=147, 73.8%) followed by A (n=41, 20.6%) and B2 (n=11, 5.5%) genotypes. Amongst the variants of BimA gene, BimA_bp_and BimA_bm_ were observed among 190 (95.4%) and 9 (4.5%) of the isolates, respectively. 190 (95.4%) study isolates harbored fhaB3 gene. None of the three LPS genotypes had a significant association with clinical forms of the disease in our study population. However, we observed a significant association between Bim A gene variants and clinical form of the disease. Association of individual genotypes/virulence genes tested with clinical form of the disease is depicted in Table 3. Using univariate analysis, we observed that carriage of BimA_bm_ gene variant had statistical significant association with neurological form of the disease (Crude OR: 7.36; 95% CI: 1.52-35.69).

**Table 3:**
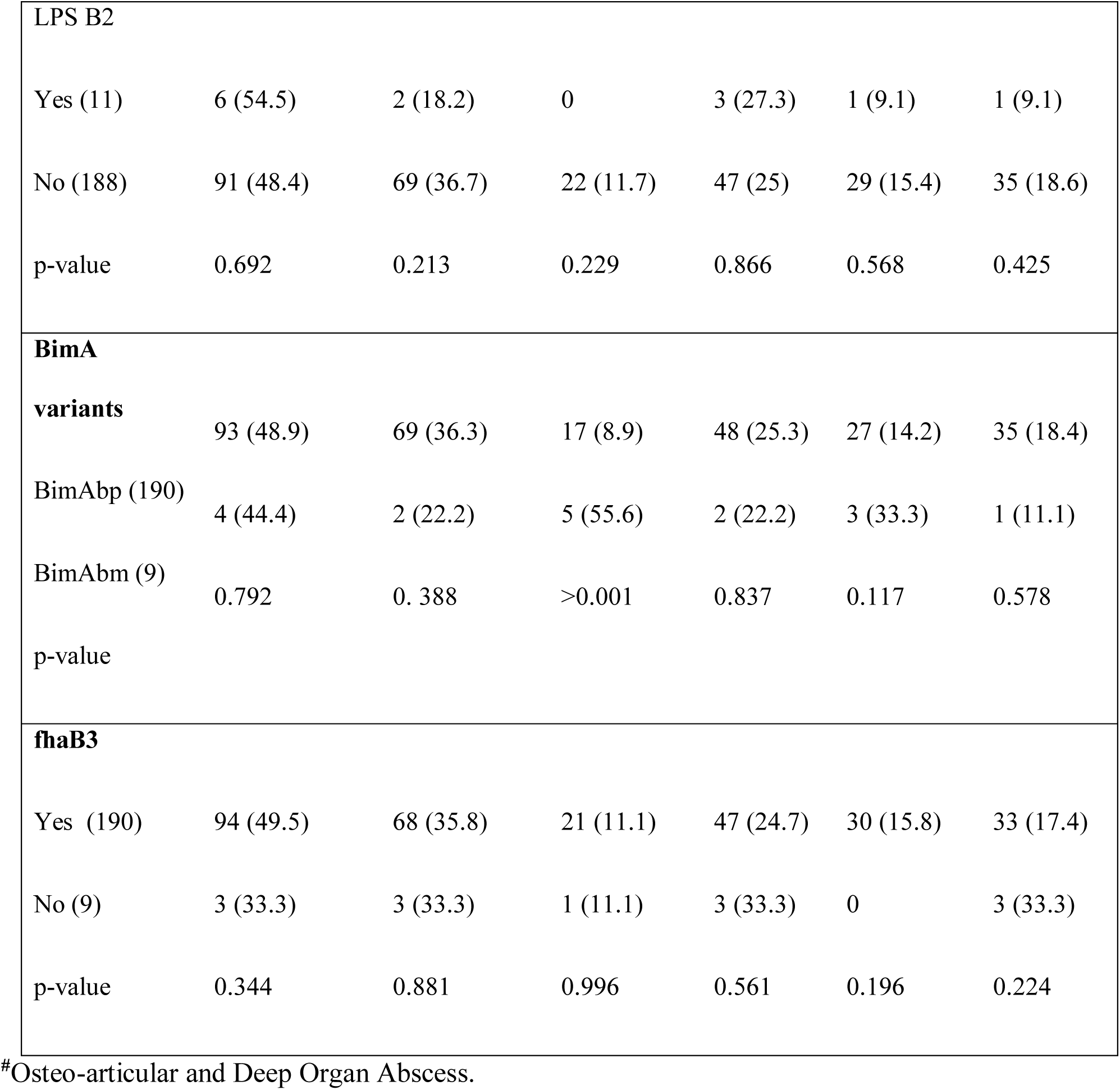
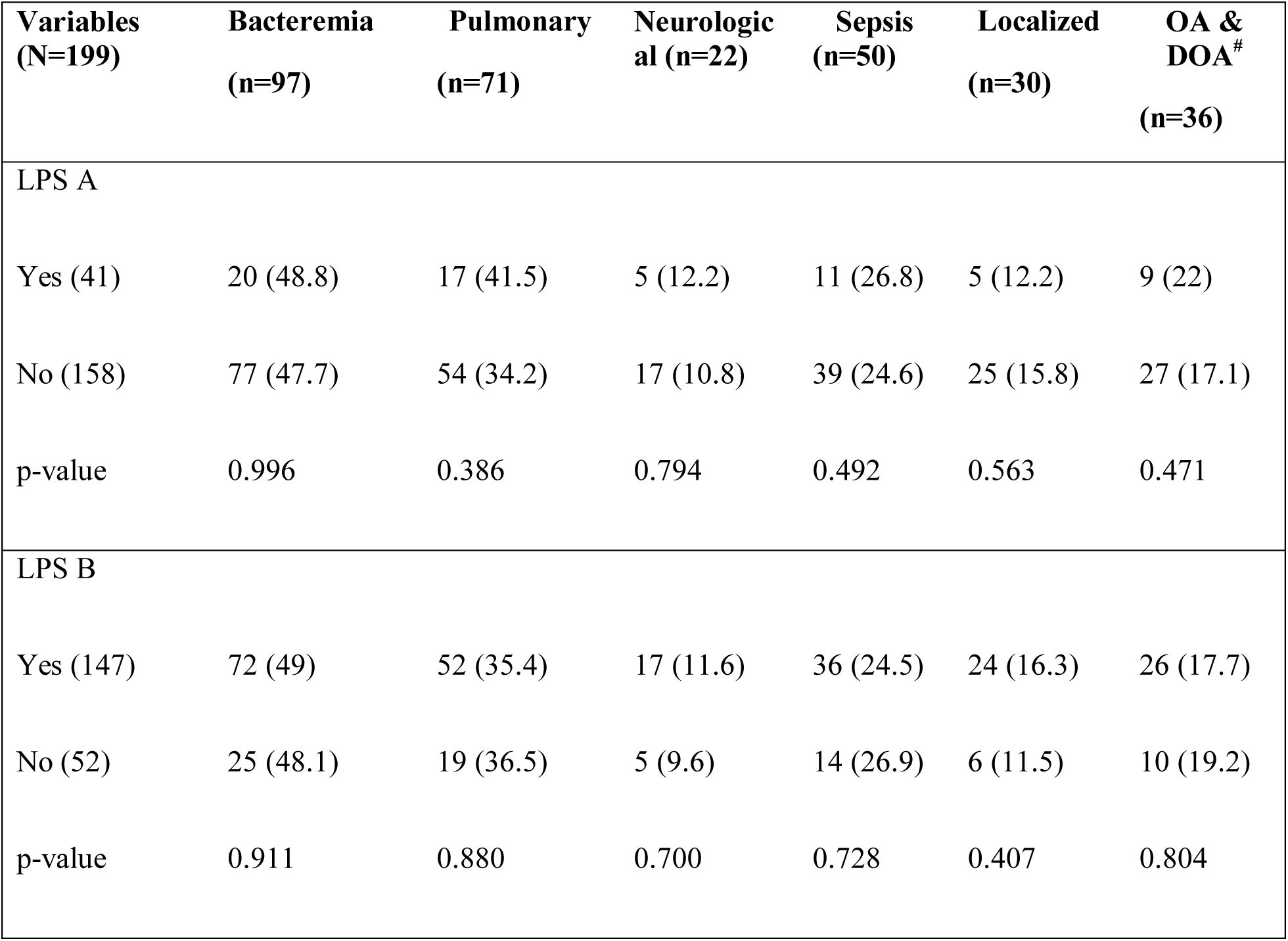
Association of lipopolysaccharide genotypes and variable virulence genes of B. pseudomallei isolates with clinical presentations in our study.

### Spatial and temporal variations of LPS genotypes in our study isolates

Spatial distribution of LPS A and B genotypes detected in the present study are depicted in **Figure 1**. All the 12 isolates obtained from patients residing in the adjacent states (8 from Kerala and 4 from Goa) belonged to the LPS-B genotype. As described above, majority (73.8%) of the *B. pseudomallei* isolates causing infection in our settings belonged to the LPS-B genotype. While, LPS-A and LPS-B genotypes were seen throughout the study period (2006-2016), isolates belonging to LPS-B2 genotype were observed only during the last three years (2014-2016). We noticed a steady decline during the years 2014-2016 in the proportion (79% in 2014, 56.6% in 2015 and 50% in 2016) of melioidosis cases due to LPS-B genotype of *B. pseudomallei* in our settings. At the same time, there was an increase in the proportion of infections due to LPS-A (16.6%, 26.6% and 33.3%) and B2 (4.1%, 16.6% and 16.6%) genotypes during the years 2014,2015 and 2016 respectively. Year-wise distribution of LPS genotypes A, B and B2 among our study isolates is depicted in **Figure 2.**

**Figure 2:**
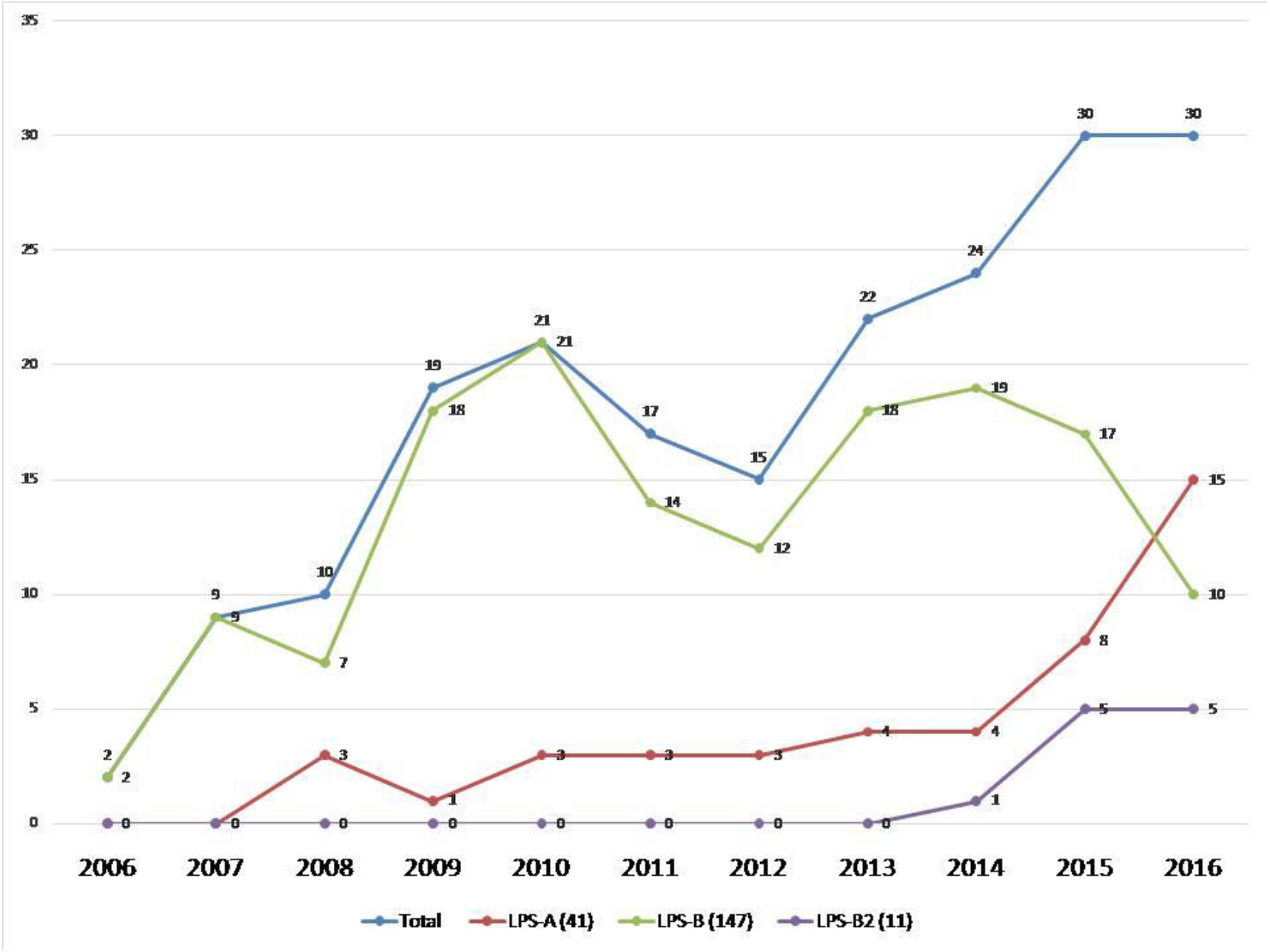
Year-wise distribution (2006-2016) of *B.pseudomallei* lipopolysaccharide genotypes causing infections in our settings.

### Influence of monthly rainfall and humidity on lipopolysaccharide diversity

Monthly rainfall and humidity data for the years 2010-2016 was plotted against time and decomposed data for trend, seasonality and residual component were compared with LPS-A and B genotypes. LPS-A genotype showed a reversal peak for trend and seasonal components of rainfall, whereas rainfall had a significant increasing effect on LPS-B genotype. There has been a rise in LPS-A genotypes observed over time, as compared to the LPS-B genotypes. On omitting the seasonal trends, the residual component showed a positive effect on LPS-B (p<0.001). The trend(p=0.04) and seasonality(p=0.01) for rainfall also showed a positive effect on LPS-B genotype, whereas seasonality only had a positive correlation (p<0.001) for LPS-A. (**Figure 3& 4**). Comparing the patterns of LPS genotypes with humidity, a positive correlation was observed between the seasonal component of humidity with LPS-A (p<0.001) and LPS-B (p=0.003). A lag period was observed in the peaks of both LPS-A and B genotypes compared to trend and seasonal component of rainfall.

**Figure 3:**
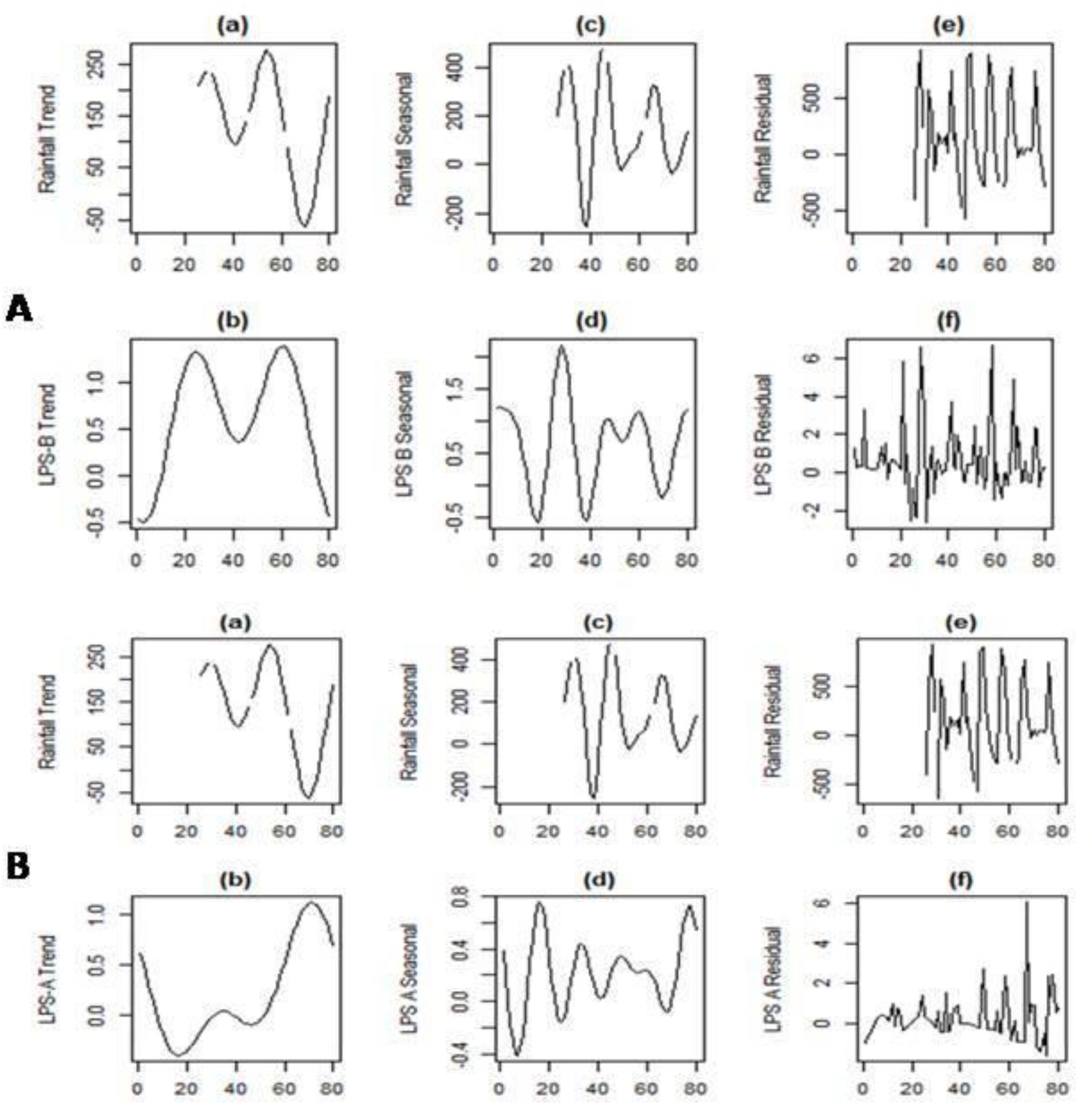
Fig A) shows reversal peaks for the trend and seasonal components of rainfall and LPS-A. B) shows an increase in LPS-B genotype with trend and seasonality. There is a lag period between the peak rainfall and the LPS-A and LPS-B genotypes.

**Figure 4:**
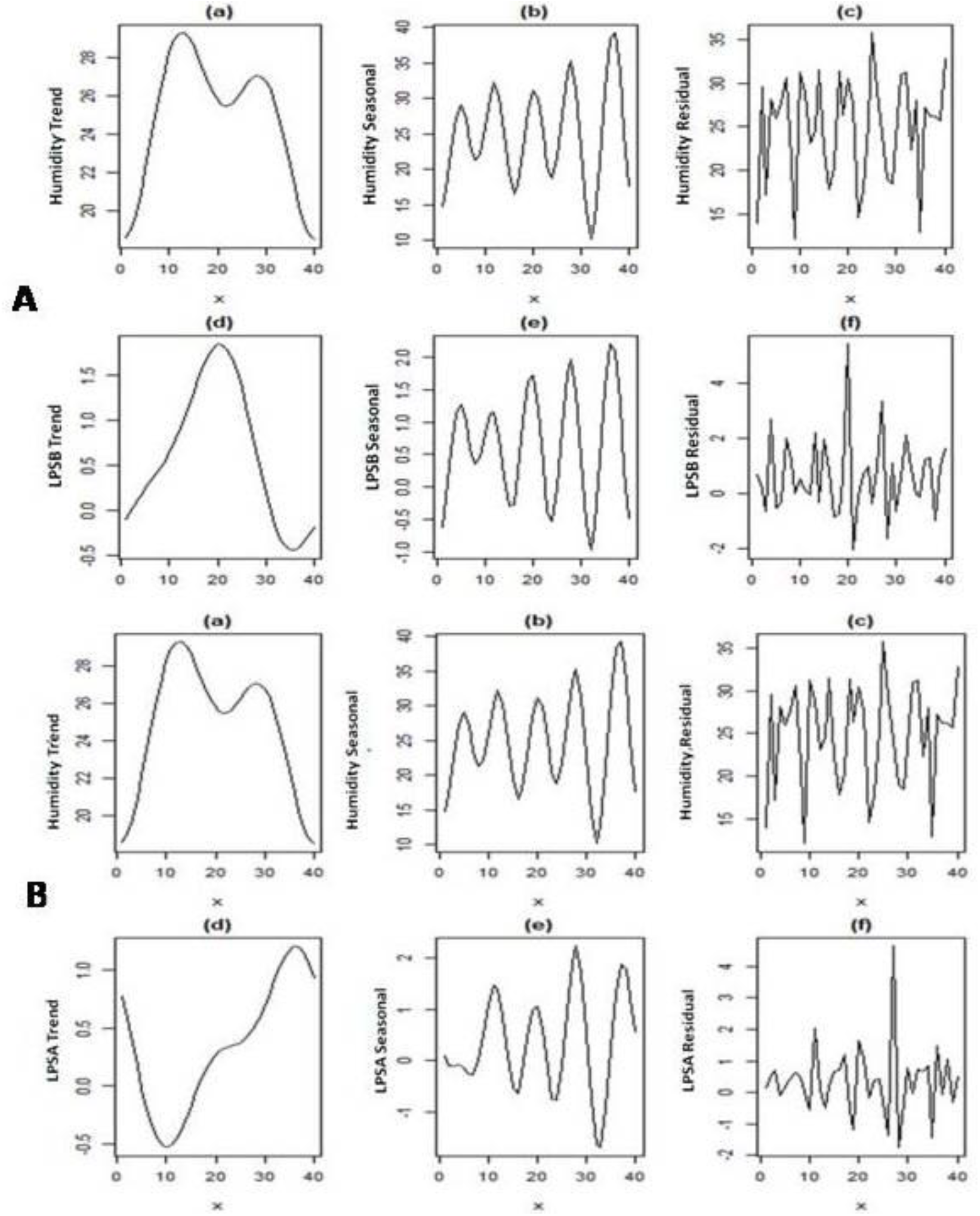
Fig A) shows reversal peaks for the trend component of humidity and LPS-A. B) shows an increase in LPS-B genotype with trend, seasonality and the residual component. There is a lag period between the peak humidity and LPS-B genotypes.

### Treatment and outcomes

Among the 199 cases, 173 (87%) patients received anti-melioidosis therapy. Of the 26 patients that did not receive pathogen-specific therapy, 17 and 9 cases were septicemic cases of melioidosis with and without bacteremia respectively. Mortality due to melioidosis was observed among 51 (25.1%) patients. None of the patients with localized form of the disease had adverse clinical outcomes. Among the 50 patients with septicemic melioidosis, 23 (46%) patients succumbed to death. Case fatality rates were 14.5% (n=25) among patients that received pathogen-specific therapy in comparison with 100 % (n=26) among those that did not receive the specific therapy.

### Host and pathogen specific determinants for clinical presentations and mortality

Renal dysfunction was an independent risk factor (after considering all the host and pathogen characteristics enlisted in **Tables 2& 3**) for both bacteremic (Adjusted OR: 3.5 (1.4-8.7), p=0.007) and septicemic (Adjusted OR: 8.75 (3.60-21.25), p>0.01) forms of the disease. Infection due to *B. pseudomallei* having BimABm variant (Adj OR: 6.7 (1.98-22.85), p=0.001) was an independent risk factor for neurological melioidosis in our study population. DM [Adjusted OR: 2.37 (95%CI: 1.19-4.74), p=0.014], septicemic melioidosis [Adjusted OR: 2.74 (95%CI: 1.31-5.85), p=0.007] and infection due to *B. pseudomallei* LPS-B genotype [Adjusted OR: 4.47 (95%CI: 1.16-12.20), p=0.003] were independent risk factors for mortality in our study population.

## Discussion

*B. pseudomallei,* the etiological agent of melioidosis demands no further negligence in view of its increasing geographical distribution, possessionof numerous virulence traits and intrinsic resistance mechanisms to several antimicrobial agents. A recent study on global evolution of *B. pseudomallei* reported geographically distinct genes/variants, conferring virulence among Australasian and Southeast Asian isolates [13]. However, there is a visible scant of whole-genomic sequencing data and/or the virulence attributes of *B. pseudomallei* isolates from regions outside Northern Australia and Thailand. Numerous studies have reported the influence of ecological factors on *B. pseudomallei* positivity in environmental niches [14,15]. We report here for the first time, the correlation of ecological factors in a geographical locality on infecting LPS genotypes of *B. pseudomallei* amongst patients.

Lipopolysaccharide of *B. pseudomallei* is an important virulence factor that facilitates the evasion of human immune responses during the early stages of infection. Monoclonal antibodies against the LPS of *B. pseudomallei* were found to reduce the severity of disease in animal models, thus implying the role of LPS as a potential vaccine candidate [1].Most intriguing finding from our study is that the majority (74%) of our patients were infected by the LPS-B genotype of *B. pseudomallei*. This observation is in contrast with the findings from Thailand (2.3%) and Australia (13.8%), where LPS-A genotype was reported to be the prevalent infecting genotype [4]. Immunological responses and disease progression among animal models administered with LPS A and B types of *B. pseudomallei* were reported previously. While, LPS-A and B are known to confer serum resistance equally and grow in the presence of 10-30% of normal human serum [16], evidence from recent experimental and animal model studies suggest that LPS-B is a more potent inducer of the pro-inflammatory cytokines and septic-shock [17]. Of the 14 isolates belonging to ST 1368 tested in the present study, 12 (85.7%) belonged to the LPS-B genotype and the rest two isolates were LPS-A genotype. This finding suggests that the LPS genotypes can vary among isolates belonging to the same sequence type and thus making it difficult to ‘brand’ a particular ST as a more or less virulent one. In the present study, we did not observe a significant association of any of the three genotypes (A, B and B2) observed with any particular clinical presentation. We observed that infection due to LPS-B genotype was an independent risk factor for mortality among our study population. However, we foresee the need for further validating this finding amongst patient populations from other geographic locations.

Expression of Burkholderia intracellular motility (Bim) A protein is crucial in the pathogenesis of the disease. Among the two variants of Bim A known, BimA_Bp_ was the only variant reported among isolates from Thailand and other South Asian countries. On the contrary, isolates from Australia were reported to have both BimA_Bp_ and BimA_Bm_ variants [4]. Among our study isolates, BimA_Bp_ variants were more commonly observed, but with no association with any particular clinical form of the disease. Presence of BimA_Bm_ was also observed in few (n=9) of our isolates and had a significant association with neurological presentations. Similar association of BimA_Bm_ variant with neurological melioidosis was reported in Australian patients and more recently in a study using mice model [4,18]. Filamentous hemagglutinin (FHA) is a surface protein of *B. pseudomallei* involved in adhesion to the host epithelial cells and formation of multinucleated giant cells [19]. fhaB3 is one of the three variable genes responsible for encoding the FHA protein [4]. Presence of fhaB3 gene was observed in almost 95% of our study isolates with no significant association with any form of the disease. fhaB3 gene positivity was reported previously among 100 % and 83% of the isolates from Thailand and Australia respectively. Further, presence of fhaB3 gene was reported in all the *B. pseudomallei* isolates obtained from Thai patients with bacteremic form of the disease and absence of fhaB3 gene was reported to have a significant association with cutaneous melioidosis among Australian patients [4].

The epidemiology of melioidosis is characterized by environmental factors where rainfall plays a key role in transmission of the disease [20]. Majority of the cases in other endemic nations are known to occur during the monsoon, when patients acquire the disease via inhalation or inoculation of the bacteria from soil and water. Occurrence of cases during the dry season, in many instances, is considered as a consequence of long latency and activation of the pathogen from latent foci [21]. In the present study, we observed that the LPS-A genotypes had a reverse correlation with rainfall. This finding suggests the possibility of *B. pseudomallei* LPS-A genotype’s persistence in dry environmental conditions, unlike the LPS-B genotypes, which had a positive correlation with rainfall and humidity. However, we did not observe a significant increase in the occurrence of infections due to the LPS-A genotypes during the dry season, to support the assumption. Further analyzing the data, a lag period was observed between the occurrence of the cases and rainfall. The observed lag period could not be determined due to lack of enough data points and unavailability of weekly rainfall data, which remains as one of the limitations of our study. Temperature did not show any correlation with the infecting genotypes in our study. This lack of correlation can be attributed to the absence of striking variations in temperature through the year, across the western coastal part of the country. On omitting the seasonal trends, LPS-B showed a correlation with the residual component, suggesting an influence of factors other than rainfall and humidity, which needs further investigations.

Among the co-morbid illnesses observed in the present study population, renal dysfunction was found to be an independent risk factor for septicemic and bacteremic forms of the disease. Diabetes mellitus (DM) was found to be an independent risk factor for mortality due to melioidosis in our settings. Strong association of DM with mortality was not surprising since 62% of the 169 patients with severe form of the disease were diabetic in our study cohort. Diabetes mellitus was reported as a risk factor for bacteremic form of the disease in patients from Thailand and northern parts of Australia [22,23]. However, we did not observe a significant association between DM and any one particular forms of the disease. Considering the high prevalence (16%) of DM amongst adult population residing in our settings [24], we foresee the need of more focused clinical studies to understand the disease progression amongst patients with controlled and uncontrolled diabetes mellitus.

In our study cohort, though not statistically significant, we observed localized melioidosis amongst a higher proportion of patients with age <18 years (28.6%) in comparison with those with age >51 years (10.5%). Skin/soft tissue infections were the common clinical presentations among cases of localized melioidosis in our settings. These infections are more likely to occur in children due to their exposure to the bacteria, while playing in water lodged fields during monsoon season. Higher frequency of localized, non-bacteremic cases was previously reported among Australian patients belonging to age group of <16 years [25]. Increased occurrence of localized form of the disease among patients of < 18 years of age can also be attributed to the lack of other predisposing factors responsible for the disease progression/dissemination Put together, the present study provides key insights regarding the disease dynamics, host and pathogen specific determinants for disease presentations and outcomes among Indian melioidosis patients.

## Notes

## Acknowledgements

The authors would like to acknowledge the contributions of the clinicians fromthe departments of Medicine, Surgery, Pediatrics, Pulmonary Medicine, Orthopedics and Neurology, Kasturba Hospital, Manipal in treating the patients with melioidosis. The authors also acknowledge Snigdha Reddy, Postgraduate student Department of Microbiology, Kasturba Medical College, Manipal for providing assistance in laboratory work. We would also like to acknowledge the National Data Center, Indian Metrological department Pune, India for providing the seasonal data for rainfall, humidity and temperature.

